# Characterization of the C4 proteins encoded by okra-infecting geminiviruses in India

**DOI:** 10.64898/2026.02.03.703481

**Authors:** Aparna Chodon, Laura Medina-Puche, Hua Wei, Gopal Pandi, Rosa Lozano-Durán

## Abstract

Okra (bhendi) is a widely cultivated food crop in warm regions of the world, with India contributing approximately 60% of global production. However, okra cultivation in India is severely constrained by viral diseases, among which infections caused by the geminiviruses bhendi yellow vein mosaic virus (BYVMV) and okra enation leaf curl virus (OELCuV), in association with their satellites, represent major limitations to crop productivity. In recent years, the geminivirus-encoded C4 protein has emerged as a key pathogenicity determinant in this viral family, with functions that include suppression of multiple layers of plant antiviral defence and induction of disease symptoms. Here, we comparatively characterize the C4 proteins of BYVMV and OELCuV by determining their targeting signals and subcellular localization, and by assessing their ability to induce developmental abnormalities and suppress the cell-to-cell spread of RNA silencing. Our results reveal that the two C4 proteins display distinct subcellular localization patterns, yet both are capable of inducing developmental alterations, likely through different mechanisms, and of suppressing the intercellular spread of RNA silencing, possibly via interaction with a common host factor. Together, these findings suggest that C4 might be a critical virulence factor in okra-infecting geminiviruses and act as a symptom determinant. The C4 proteins encoded by BYVMV and OELCuV therefore emerge as promising targets for the development of antiviral management strategies in okra.

## INTRODUCTION

Okra or bhendi (*Abelmoschus esculentus*) is a food crop cultivated in tropical, subtropical, and warm or temperate regions of the globe; it is a staple vegetable in the diets of populations in South Asia, sub-Saharan Africa, the Middle East, and parts of the Caribbean, providing an affordable source of dietary fibre, vitamins, and micronutrients. Global demand for okra is moreover steadily increasing, driven not only by population growth in producing regions, but also by expanding international trade and rising interest in plant-based foods.

India is the world’s largest okra producer, contributing approximately 60% of the global harvest (FAOSTAT). However, okra cultivation in India is threatened by viral diseases, many of which are caused by members of the family *Geminiviridae*. Geminiviruses are insect-transmitted plant-infecting viruses with circular, single-stranded DNA genomes, responsible for devastating crops diseases worldwide. The two main geminiviral complexes constraining okra production in India are bhendi yellow vein mosaic virus (BYVMV) (*Begomovirus abelmoschusflavi*) and okra enation leaf curl virus (OELCuV) (*Begomovirus abelmoschusenation*), in association with their respective viral satellites. BYVMV, together with its associated betasatellite, is the causal agent of yellow vein mosaic disease (1); OELCuV likely leads to enation leaf curl disease, when co-infecting either with its associated alphasatellite or with BYVMV betasatellite (2,3). Mixed infections of BYVMV and OELCuV in okra has also been reported (4). Both viruses belong to the genus Begomovirus (family *Geminiviridae*) and present the typical monopartite genome organization, with seven protein-coding genes described so far and an intergenic region containing the origin of replication as well as promoters, while the satellites are believed to encode one protein each (Figure 1a). In the experimental host *Nicotiana benthamiana*, full development of symptoms requires the presence of the satellites (Figure 1b) (1,3).

**Figure 1.**
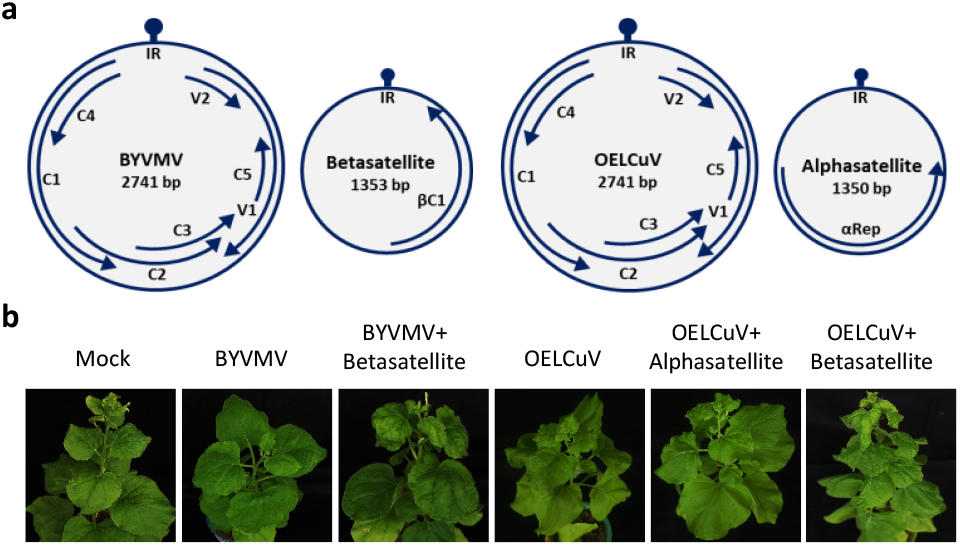
Genomic organization of okra-infecting geminiviruses and symptoms caused by these viruses in *Nicotiana benthamiana*. (a) Genome organization of bhendi yellow vein mosaic virus (BYVMV) and its associated betasatellite, and okra enation leaf curl virus (OELCuV) and its associated alphasatellite. The size of the DNA molecules (in bp) is shown. Genes are depicted by arrows; gene names are indicated. IR: intergenic region. The lollipops indicate the stem loop at the origin of replication. (b) *Nicotiana benthamiana* plants mock-inoculated or inoculated with infectious clones of BYVMV, BYVMV together with its associated betasatellite, OELCuV, and OELCuV together with its associated alphasatellite or with the BYVMV betasatellite at 28 days post-inoculation (dpi)

Despite the economic and humanitarian relevance of the viral diseases infecting okra, the underlying molecular mechanisms remain largely elusive. In recent years, the begomovirus-encoded C4 protein has emerged as a crucial pathogenicity factor in this clade of viruses, displaying the ability to suppress different layers of the plant antiviral defence and to act as a symptom determinant, frequently leading to severe developmental alterations in infected plants (5). The C4 protein from BYVMV has been shown to be required for systemic infection but not viral replication, to act as a suppressor of post-transcriptional gene silencing (PTGS) as well as methylation-mediated transcriptional gene silencing (TGS) (6–8), and to cause developmental alterations when transgenically expressed in *N. benthamiana* (Gopal et al., 2007); whether these properties are shared by the C4 protein from OELCuV is at present unknown. It is also unclear whether either of these C4 proteins impacts symptom development during infection, as shown for geminiviral species infecting other hosts (5).

In this work, we perform a comparative characterization of the C4 proteins encoded by the two main okra-infecting geminiviruses in India, BYVMV and OELCuV, capitalizing of the existing body of knowledge about this geminiviral protein. With the aim to gain insight into the functional contribution of these proteins to the infection, we begin by analysing the targeting signals and known domains present in their sequence, investigate their subcellular localization, and assess their ability to alter plant development and to suppress the cell-to-cell movement of RNA silencing, described for the C4 proteins from other geminiviruses (Medina-Puche et al., 2021). Next, we evaluate the capacity of these viral proteins to interact with known targets of C4 proteins encoded by other geminiviruses and responsible for the aforementioned activities. Our results indicate that each of these two C4 proteins contains different motifs/domains and shows specific localization patterns; however, both can induce developmental alterations, probably through distinct mechanisms, and suppress the intercellular spread of RNA silencing, likely through targeting the same plant protein. We propose that C4 is likely a critical virulence factor for BYVMV and OELCuV, and that it might be a symptom determinant in okra. The C4 proteins encoded by these viruses therefore emerge as potential promising targets for the development of antiviral management strategies in this crop.

## RESULTS

### The C4 proteins encoded by BYVMV and OELCuV have distinct features and subcellular localization patterns

Even though BYVMV and OELCuV both infect okra in a common geographical area, their encoded C4 proteins show only 40% sequence identity, suggesting that they may diverge in function (Figure 2a). C4 proteins from different geminiviruses frequently display combinations of two targeting signals, namely a myristoylation site that tethers them to membranes and a chloroplast transit peptide (cTP) that targets them to this organelle (5); the presence/absence of these signals likely influences the distribution of the protein pool within the host cell. In order to gain insight into the subcellular localization and potential functions of the C4 proteins encoded by the okra-infecting BYVMV and OELCuV, we first predicted the presence of myristoylation sites and cTPs in their sequence. A potential N-myristoylation site was identified in BYVMV C4, but not in OELCuV C4, while none of these proteins contained a predicted cTP. OELCuV C4 was found to harbour a CCL-like domain, recently described in the C4 protein encoded by tomato yellow leaf curl virus (TYLCV) (9) (Figure 2b, c).

**Figure 2.**
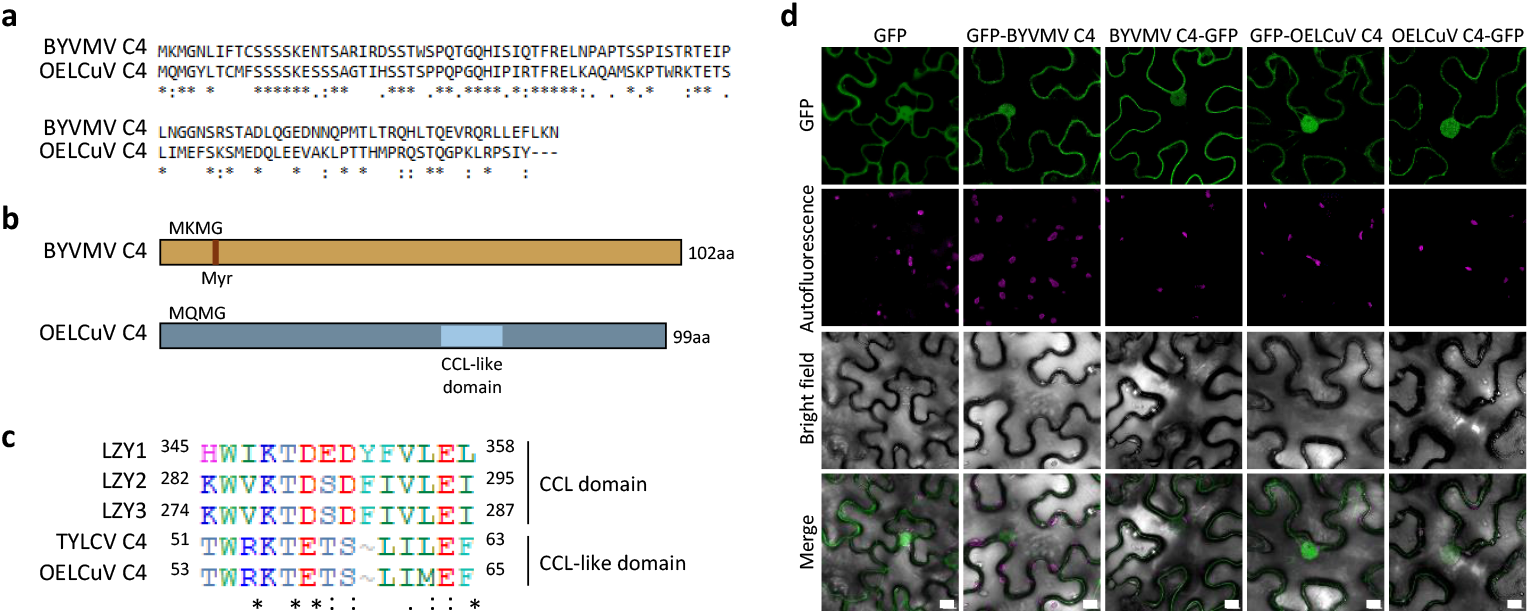
Sequence comparison, predicted features, and subcellular localization of GFP-fused BYVMV and OELCuV C4 proteins in *N. benthamiana* epidermal cells. (a) Pairwise alignment showing sequence identity between BYVMV and OELCuV C4 sequences using Needle Pairwise Alignment tool. (b) Schematic representation of the predicted features on BYVMV or OELCuV C4; size (in aa) is indicated. Myr: myristoylation site, as predicted by Expasy Myristoylator. (c) Alignment of amino acid sequences of LZY1, LZY2, and LZY3 in the region containing the CCL domain, and the CCL-like domain in TYLCV C4 and OELCuV C4. Numbers indicate the amino acid position in the corresponding proteins. Asterisks denote identical residues, while colons denote similar residues. (d) Subcellular localization of GFP-fused BYVMV or OELCuV C4 proteins in *N. benthamiana* leaves infiltrated with *A. tumefaciens* carrying constructs to express free GFP, GFP-C4, or C4-GFP. Samples were observed under the confocal microscope at 2 days post-inoculation (dpi). Scale bar: 10 μm.

Subcellular localization experiments following transient expression of GFP-fused proteins in epidermal pavement cells of *N. benthamiana* leaves revealed an apparent plasma membrane association of BYVMV C4-GFP, while GFP-C4, in which the fluorescent protein would potentially interfere with N-myristoylation of C4, seems distributed in the nucleus and the cytoplasm, with a pattern indistinguishable from that of free GFP (Figure 2d). A similar localization pattern is displayed by both OELCuV GFP-C4 and C4-GFP (Figure 2d). The plasma membrane localization of BYVMV C4-GFP was confirmed by its co-localization with the plasma membrane marker LTI6B (10) (Figure 3a; Figure S1a). Interestingly, even though no clear plasma membrane association could be detected for C4 from OELCuV, this protein could be found in plasmodesmata, as indicated by its co-localization with the plasmodesmata marker PDLP5 (11) (Figure 3b); a small fraction of this C4 protein can also be observed in chloroplasts (Figure 2d). Both C4 proteins could also be found in the nucleus, co-localizing with H2B (12) (Figure S1b).

**Figure 3.**
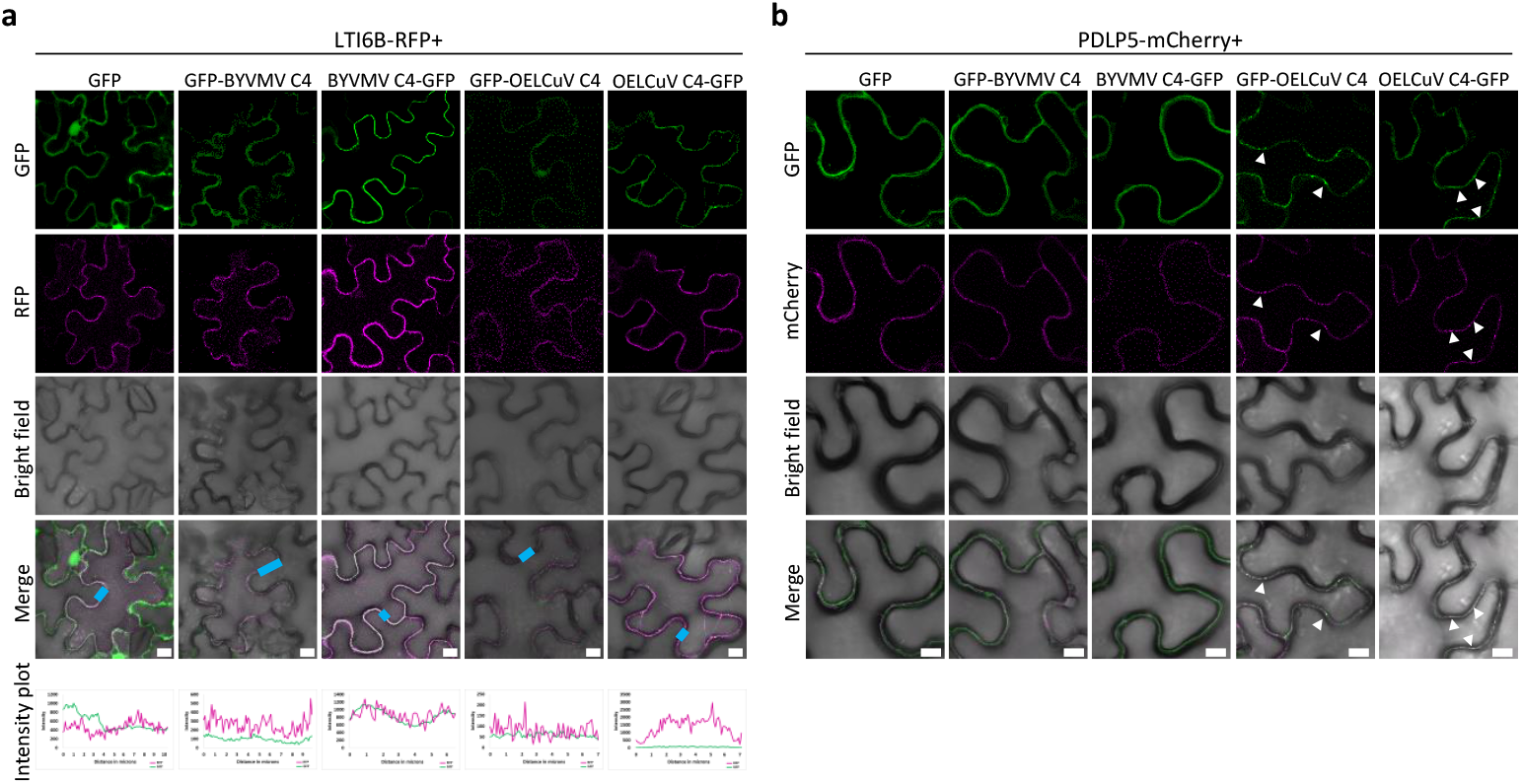
Co-localization analysis of GFP-fused BYVMV and OELCuV C4 proteins in *N. benthamiana* epidermal cells. (a) Co-localization analysis of GFP-fused C4 proteins and the plasma membrane marker LTI6B-RFP in *N. benthamiana* leaves co-infiltrated with *A. tumefaciens* carrying constructs to express the marker, GFP-C4, C4-GFP, or free GFP as a negative control. Colocalization between the GFP and RFP channels is analyzed with line scan intensity plots (indicated by the blue line in the merge image). Maximum projection images of z-stacks are shown in Figure S1A. (b) Co-localization analysis of GFP-fused C4 proteins and the plasmodesmata marker PDLP5-mCherry in *N. benthamiana* leaves co-infiltrated with *A. tumefaciens* carrying constructs the marker, GFP-C4, C4-GFP, or free GFP as a negative control. Spots in which proteins are co-localized are indicated with white arrowheads. Samples were observed under the confocal microscope at 2 days post-inoculation (dpi). Scale bar: 10 μm.

Taken together, our results indicate that the C4 proteins encoded by these two okra-infecting geminiviruses differ in sequence, motifs, and subcellular localization, raising the possibility that they may intersect differently with the host cell machinery.

### The C4 proteins encoded by BYVMV and OELCuV induce developmental alterations resembling symptoms when expressed *in planta* and show a differential interaction with the known symptom-related targets NbSKeta and RLD3

C4 proteins from multiple geminiviruses have been shown to act as symptom determinants, causing the plant developmental alterations recognized as symptoms during infection (5,13). To assess whether the C4 proteins from BYVMV and OELCuV interfere with plant development, we first expressed them systemically in *N. benthamiana* from a potato virus X (PVX)-based vector. Interestingly, both C4 proteins caused leaf distortion, with newly formed leaves being smaller in size and showing changes in coloration when compared to control plants (inoculated with an empty PVX-derived vector or with a vector containing a fragment of the GFP-coding gene with a size similar to that of the C4-coding genes) (Figure 4a, b). To test the effect of these C4 proteins in a different plant species, we next generated transgenic *Arabidopsis thaliana* (hereafter referred to as Arabidopsis) plants expressing them from a constitutive 35S promoter (Figure 4c, d; Figure S2). As shown in Figure 4c and d, both C4 proteins cause severe developmental malformations in this species, with transgenic plants displaying smaller size and twisted and curly leaves.

**Figure 4.**
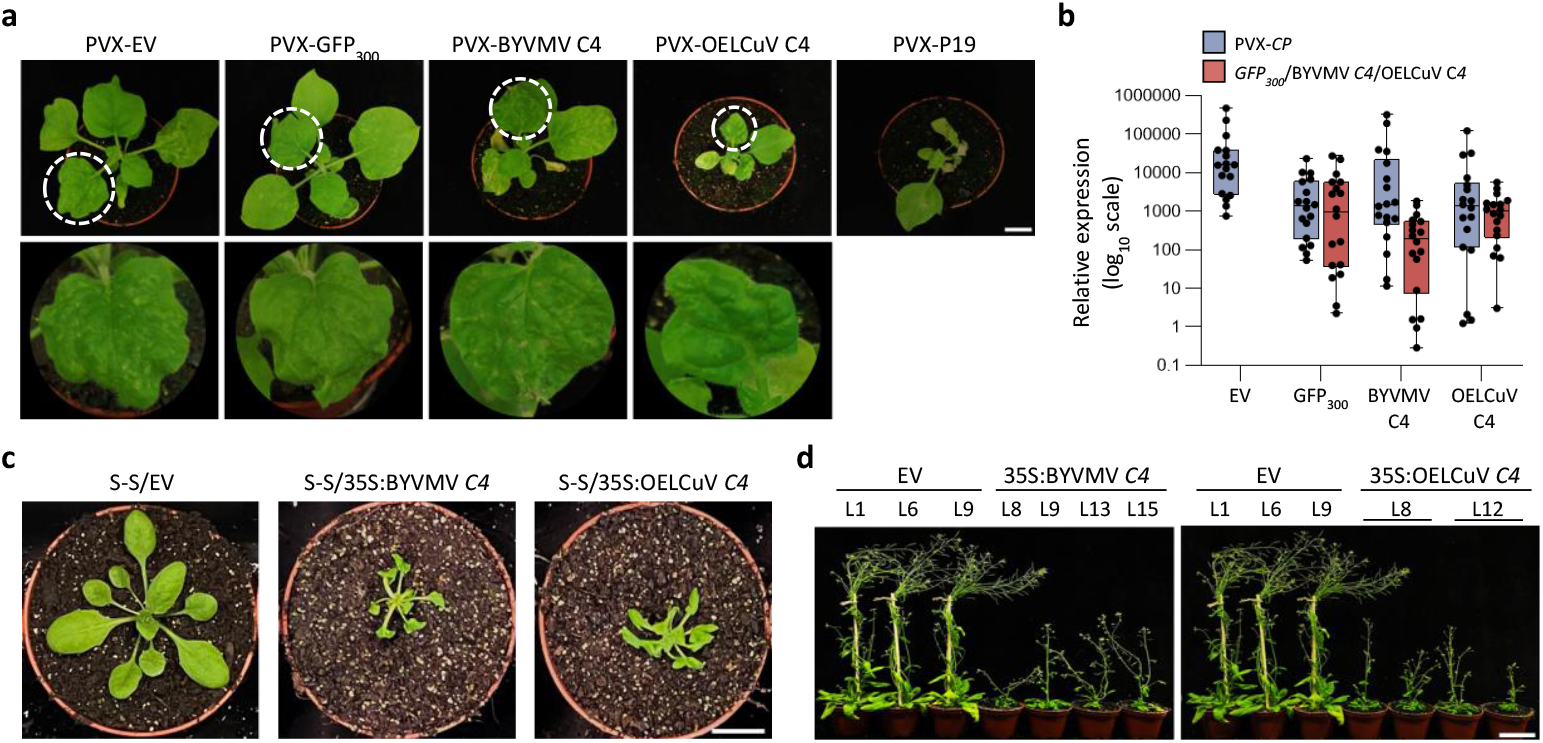
C4 proteins from BYVMV and OELCuV cause developmental alterations. (a) *N. benthamiana* plants infected with a PVX-based vector expressing BYVMV C4, OELCuV C4, P19, or containing a 300-bp GFP fragment.17-day-old plants were taken for the infection. Pictures were taken at 9 days post-inoculation (dpi). PVX-EV (empty vector) or PVX-GFP_300_ and PVX-P19 used as negative and positive controls, respectively. Scale bar: 3 cm. (b) Accumulation of PVX CP, GFP_300,_ BYVMV C4, and OELCuV C4 transcripts in plants in (a) relative to actin mRNA, as measured by qRT-PCR. Two newly emerging leaves per plant were used for the quantification. The infection was performed in three independent replicates with six plants in each replicate and the boxes represent the aggregate data obtained from all the three replicates. (c) Four-week-old transgenic *Arabidopsis thaliana* plants expressing BVVMV C4 or OELCuV C4 under a 35S promoter (35S:BYVMV C4 or 35S:OELCuV C4) or transformed with the empty vector (EV) as control, grown in long-day conditions. One representative line is shown per construct. Scale bar: 2 cm. (d) Seven-week-old transgenic *A. thaliana* plants expressing BVVMV C4 or OELCuV C4 under a 35S promoter (35S:BYVMV C4 or 35S:OELCuV C4) or transformed with the empty vector (EV) as control, grown in long-day conditions; independent lines (LX) are shown. The EV-transformed plants shown in both panels for comparison are the same. Scale bar: 6 cm.

Two different mechanisms have been so far proposed to explain the capacity of C4 to cause developmental alterations in plants. C4 from tomato leaf curl Yunnan virus (TLCYnV) interacts with the kinase NbSKη and re-localizes it from the nucleus to the plasma membrane, ultimately triggering additional cell divisions (14,15). C4 from TYLCV, which does not interact with NbSKη, binds members of the plant-specific RCC1-Like Domain (RLD) protein family and recruits them to the cell periphery, possibly impairing their function in the establishment of cell polarity and consequently polar auxin transport (9). The interaction of C4 with NbSKη or with RLD proteins has been shown to require defined protein motifs, namely a minidomain in TLCYnV C4 and the CCL-like motif in TYLCV C4 (9,16); while none of the C4 proteins from these okra-infecting geminiviruses seems to contain an obvious NbSKη-interacting minidomain, that encoded by OELCuV harbours a CCL-like domain, as previously mentioned (Figure 2b, c).

To gauge if the potential symptom determination activity of these C4 proteins might rely on one of the mechanisms already described, we decided to test their interaction with NbSKη and RLD3 in co-immunoprecipitation (co-IP), bimolecular fluorescence complementation (BiFC), and yeast two-hybrid (Y2H) assays. Our results consistently indicate that BYVMV C4, but not OELCuV C4, interacts with NbSKη (Figure 5a-c; Figure S3), while the opposite is true for RLD3, as expected from the presence of a CCL-like motif in OELCuV C4 (Figure 5d-f; Figure S3). These findings suggest that the two C4 proteins might induce symptom development through distinct mechanisms, and that BYVMV C4 must contain a different NbSKη-interacting domain to that described in TLCYnV C4.

**Figure 5.**
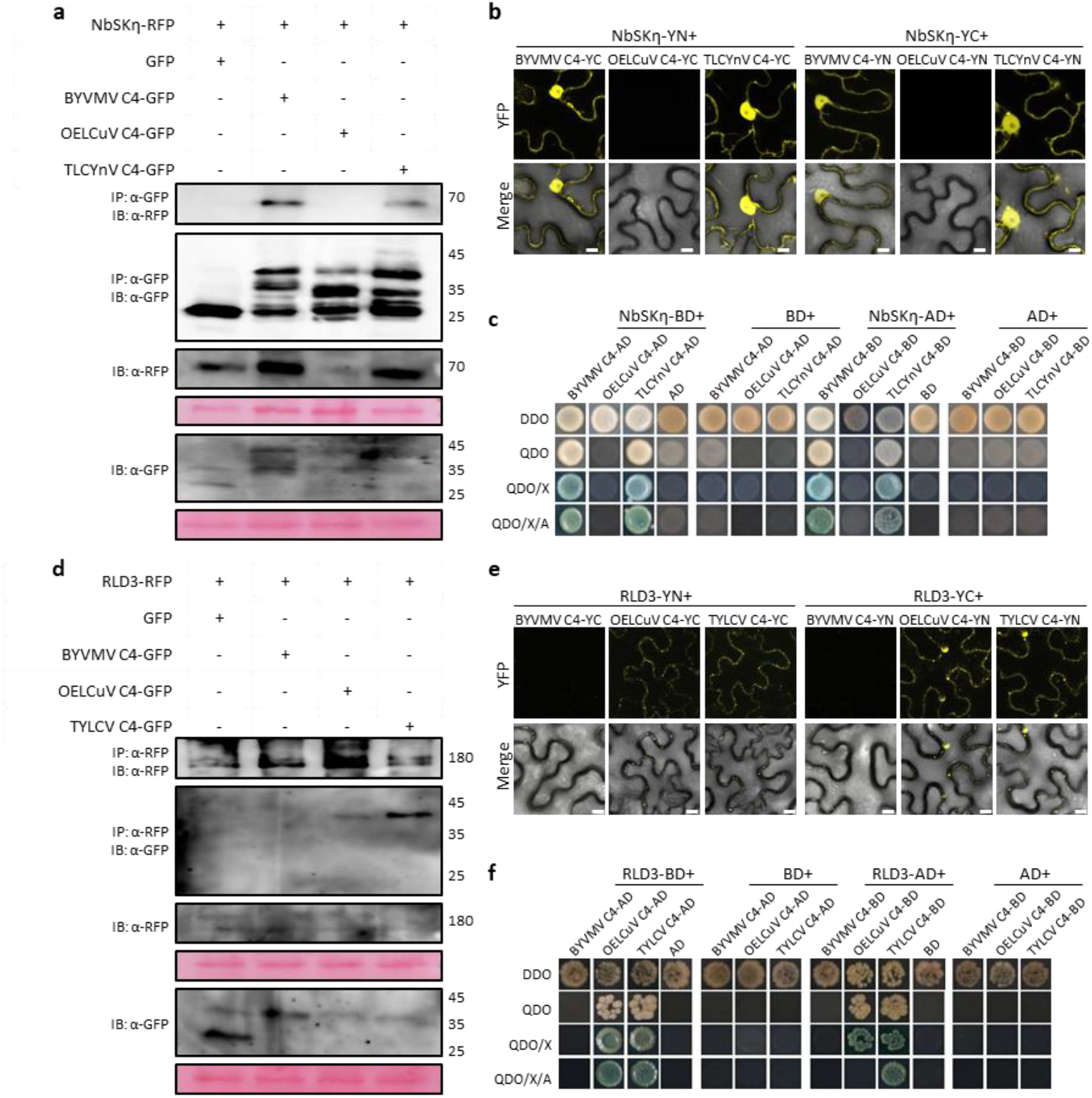
C4 proteins from BYVMV and OELCuV interact with NbSKη and RLD3, respectively. (a) Co-immunoprecipitation of NbSKη-RFP with GFP-fused BYVMV or OELCuV C4 (C4-GFP) upon transient co-expression in *N. benthamiana* leaves. Free GFP and tomato leaf curl Yunnan virus (TLCYnV) C4-GFP are used as negative and positive controls, respectively. Numbers on the right indicate molecular weight (in kDa). IP: immunoprecipitation; IB: immunoblot. Additional independent replicates are shown in Supplementary figure 3. (b) Interaction between BYVMV or OELCuV C4 and NbSKη by bimolecular fluorescence complementation (BiFC) upon transient co-expression of fusion proteins in *N. benthamiana* leaves. TLCYnV C4 is used as positive control. Images were taken at 2 days post-inoculation (dpi). Scale bar: 10 μm. (c) Interaction between BYVMV or OELCuV C4 and NbSKη by yeast two-hybrid assay. DDO (double dropout medium): SD/−Leu/−Trp. QDO (quadruple dropout medium): SD/− Ade/− His/− Leu/− Trp. QDO/X: QDO supplemented with X-α-gal (X). QDO/X/AbA: QDO supplemented with X-α-gal (X) and aureobasidin A (AbA). AD: activation domain; BD: binding domain. (d) Co-immunoprecipitation of RLD3-RFP with GFP-fused BYVMV or OELCuV C4 (C4-GFP) upon transient co-expression in *N. benthamiana* leaves. Free GFP and tomato yellow leaf curl virus (TYLCV) C4-GFP are used as negative and positive controls, respectively. Numbers on the right indicate molecular weight (in kDa). IP: immunoprecipitation; IB: immunoblot. Additional independent replicates are shown in Supplementary figure 3. (e) Interaction between BYVMV or OELCuV C4 and RLD3 by BiFC upon transient co-expression of fusion proteins in *N. benthamiana* leaves. TYLCV C4 is used as positive control. Images were taken at 2 dpi. Scale bar: 10 μm. (f) Interaction between BYVMV or OELCuV C4 and RLD3 by yeast two-hybrid assay.

### The C4 proteins encoded by BYVMV and OELCuV suppress the cell-to-cell movement of RNA silencing in the SUC-SUL reporter Arabidopsis line and interact with BAM1

The C4 proteins from two different geminiviruses, TYLCV and mungbean yellow mosaic virus (MYMV), have been shown to localize to plasmodesmata and interfere with the cell-to-cell movement of RNA silencing, considered the main antiviral defence mechanism in plants (17,18). Both proteins interact with the receptor kinase BARELY ANY MERISTEM 1 (BAM1) in plasmodesmata, an interaction that has been proposed to mediate the inhibition of RNA silencing movement (17,18). The ability to interact with BAM1 is conserved in the C4 protein from at least one more geminivirus, tomato leaf curl Guangdong virus (ToLCGdV) (19).

To evaluate the capacity of BYVMV C4 and OELCuV C4 to suppress the intercellular movement of RNA silencing, we generated transgenic SUC:SUL (S-S) reporter plants (20) expressing these viral proteins (Figure 6a-c). The SUC:SUL plants contain a construct to produce an inverted repeat of the endogenous *SULFUR* (*SUL*) gene (At4g18480), encoding a magnesium chelatase required for chlorophyl biosynthesis, in phloem companion cells, which leads to the production of small interfering (si)RNAs against *SUL* and silencing of the gene. RNA silencing can move to 10–15 cells beyond companion cells, producing a bleaching phenotype around the vasculature. Visual assessment of the extension of the bleaching indicates reduced spread in the transgenic plants expressing C4 (Figure 6b), an observation confirmed by quantification of the bleaching percentage through image analysis (Figure 6d) and supporting the notion that both C4 proteins interfere with the cell-to-cell movement of RNA silencing in this reporter system.

**Figure 6.**
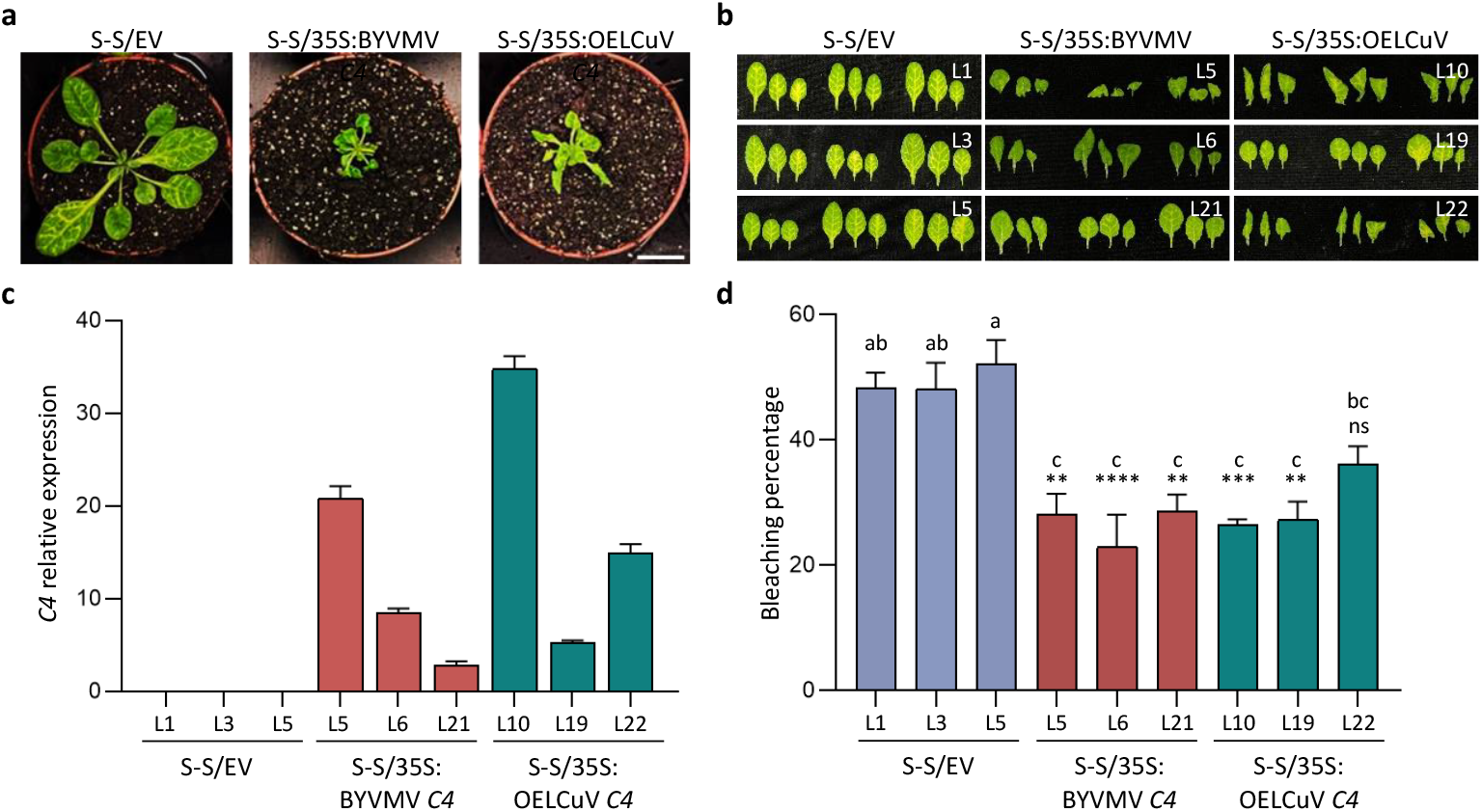
C4 proteins from BYVMV and OELCuV suppress the cell-to-cell movement of RNA interference. (a) Four-week-old SUC:*SUL* transgenic *Arabidopsis thaliana* plants expressing BVVMV or OELCuV C4 under a 35S promoter (S-S/35S:BYVMV C4 or S-S/35S:OELCuV C4) or transformed with the empty vector (S-S/EV) grown in long-day conditions. Scale bar: 2 cm. (b) Leaves of four-week-old transgenic S-S/35S:BYVMV C4, S-S/35S:OELCuV C4, or S-S/EV plants. Each set of three leaves comes from one T2 plant. Three plants have been used here to represent three independent lines (LX) for each construct. Scale bar: 1 cm. (c) Accumulation of C4 transcripts in transgenic plants relative to actin mRNA, as measured by qRT-PCR. Each bar represents the mean of five 2-week-old seedlings from each independent line. Error bars represent SEM. (d) Quantification of the bleaching percentage of the leaves from (b). Error bars represent SEM. Bars with same letter are not significantly different (P=0.05) according to Dunnet’s multiple comparison test. Asterisks indicate a statistically significant difference compared to S-S/EV. **: p < 0.01; ***: p < 0.001; ****: p < 0.0001; ns: not significant.

Next, we tested the capacity of BYVMV C4 and OELCuV C4 to interact with BAM1 in co-IP, BiFC, and Y2H assays. In all experiments, both C4 proteins interacted with BAM1 (Figure 7; Figure S3), raising the idea that targeting of this plant protein might underlie the ability of the virulence factors to suppress the intercellular spread of RNA silencing.

**Figure 7.**
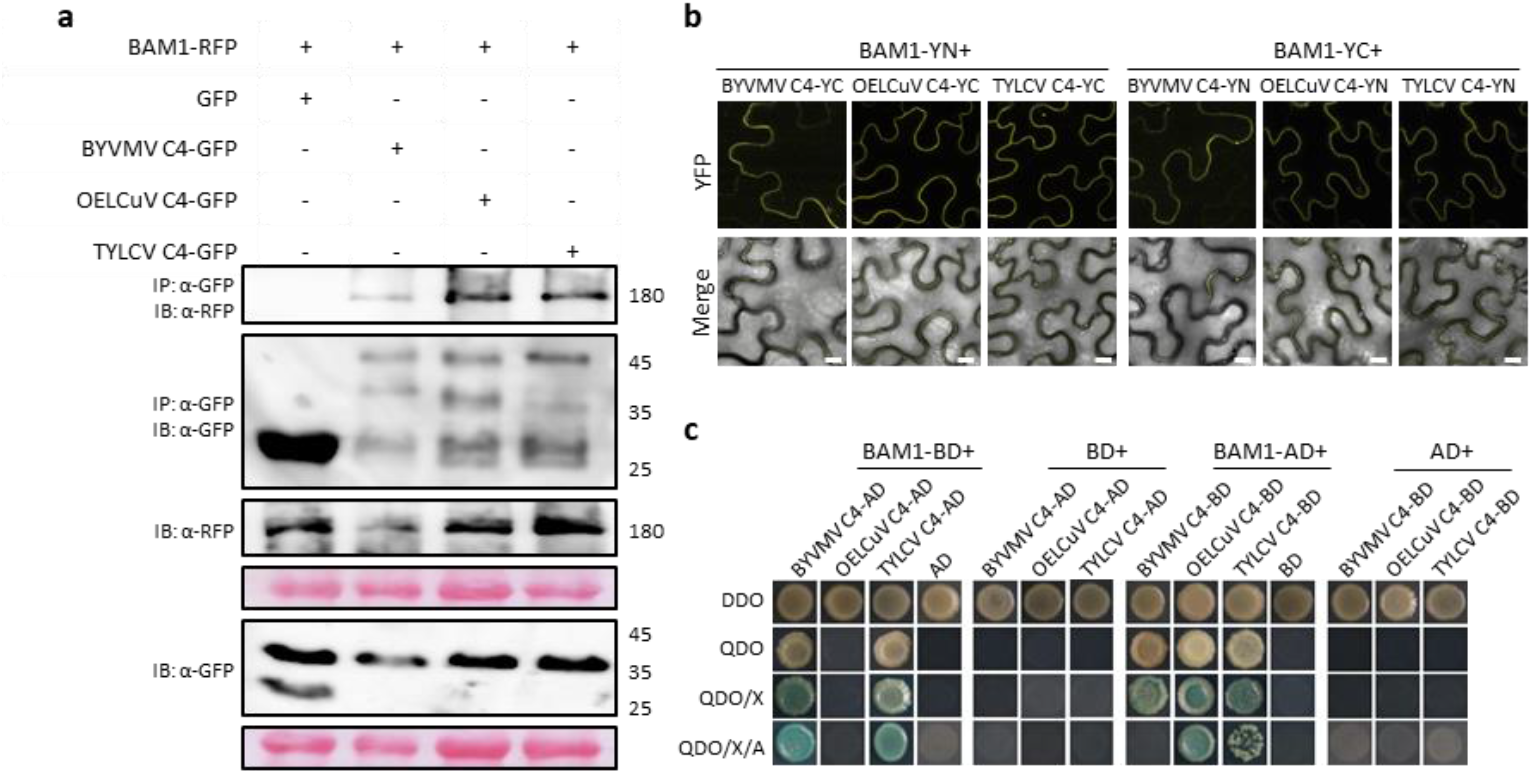
C4 proteins from BYVMV and OELCuV interact with BAM1. (a) Co-immunoprecipitation of AtBAM1-RFP with GFP-fused BYVMV or OELCuV C4 (C4-GFP) upon transient co-expression in *N. benthamiana* leaves. Free GFP and tomato yellow leaf curl virus (TYLCV) C4-GFP are used as negative and positive controls, respectively. Numbers on the right indicate molecular weight (in kDa). IP: immunoprecipitation; IB: immunoblot. Additional independent replicates are shown in Supplementary figure 3. (b) Interaction between BYVMV or OELCuV C4 and AtBAM1 by bimolecular fluorescence complementation (BiFC) upon transient co-expression of fusion proteins in *N. benthamiana* leaves. TYLCV C4 is used as positive control. Images were taken at 2 days post-inoculation (dpi). Scale bar: 10 μm. (c) Interaction between BYVMV or OELCuV C4 and the kinase domain of AtBAM1 by yeast two-hybrid assay. DDO (double dropout medium): SD/−Leu/−Trp. QDO (quadruple dropout medium): SD/−Ade/−His/−Leu/−Trp. QDO/X: QDO supplemented with X-α-gal (X). QDO/X/AbA: QDO supplemented with X-α-gal (X) and aureobasidin A (AbA).

## DISCUSSION

In spite of the growing importance of okra cultivation, our understanding of the diseases that threaten its production is still limited. This is the case for the two main viruses affecting okra in India, BYVMV and OELCuV, of which the infection mechanisms have only recently begun to be dissected. In this work, we take advantage of the previous knowledge on the geminivirus-encoded C4 protein, a well-studied crucial virulence factor in the genus begomovirus, to perform a comparative study with the aim to shed light on the properties and potential functions of the positional homologues in BYVMV and OELCuV.

Our analyses show that the C4 proteins from these two okra-infecting geminiviruses differ in motifs, localization pattern, interactors, and their effect on plant development. Both of them, however, can suppress the intercellular spread of RNA silencing in the SUC:SUL reporter system, and interact with the plant BAM1. These results underscore the potential multifunctionality of the C4 proteins encoded by these okra-infecting geminiviruses, as previously shown for others: while at least BYVMV C4 functions as a PTGS and TGS suppressor, both BYVMV C4 and OELCuV C4 can inhibit the cell-to-cell movement of RNA silencing and interfere with plant developmental programs, probably contributing to symptom development in the context of the infection (Figure 8). Additional functions of these proteins are likely, considering what has been described for other C4 proteins, and the fact that both of them also localize to the nucleus (Figure S1b).

**Figure 8.**
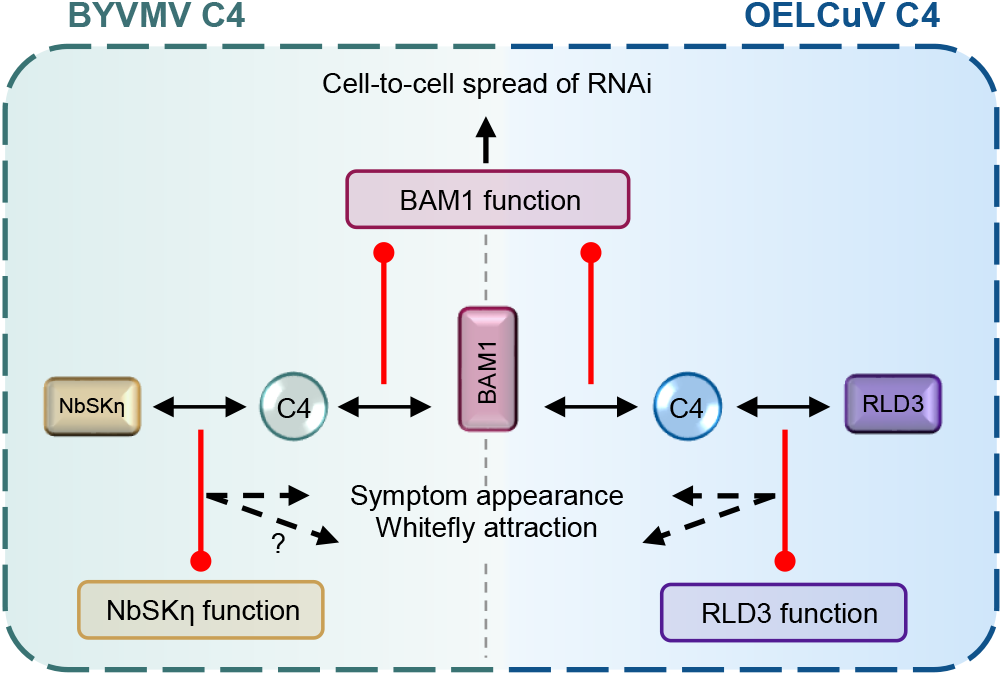
C4 proteins encoded by the okra-infecting geminiviruses BYVMV and OELCuV likely function as symptom determinants and suppressors of the cell-to-cell movement of RNA silencing. OELCuV C4 and BYVMV C4 interact with RLD3 and NbSKη, respectively, likely leading to symptom development, as described for other geminiviruses. Both C4 proteins interact with the receptor kinase BAM1and suppress the intercellular spread of RNA silencing.

While expression of either BYVMV C4 or OELCuV C4 results in developmental alterations in *N. benthamiana* and Arabidopsis, the phenotypes caused by each of them are not identical (Figure 4). This might be at least partially explained by their divergent interactions with the two C4-targeted symptom-related plant proteins described so far: while BYVMV C4 interacts with NbSKη but not RLD3, OELCuV C4 interacts with RLD3 only (Figure 5). Although the involvement of these interactions in symptom development in okra remains to be tested, their specificity supports the idea that C4 proteins might generally use one of two different strategies to interfere with development. Since BYVMV and OELCuV have been found in mixed infections in okra (4), it is possible that when co-existing in the host both C4 proteins might simultaneously interfere with plant development through these two mechanisms, potentially exacerbating symptom severity.

Intriguingly, only mild symptoms were described in *N. benthamiana* infected with BYVMV or OELCuV in the absence of their associated satellites, despite C4 being presumably expressed in these plants (Figure 1; (3,6)). Several non-mutually exclusive hypotheses could be raised to justify this observation: 1) C4 might have host-specific effects, which could explain the milder alterations caused by protein expression in *N. benthamiana* compared to Arabidopsis; 2) Viral load in the experimental host *N. benthamiana* may not be high enough to induce symptoms; 3) Expression of C4 from the virus’ genomic context might require factors not available in *N. benthamiana*, and therefore only occur when the viral gene is heterologously expressed (e.g. from a PVX-based vector or a 35S promoter). Regardless, the fact that both C4 proteins can produce developmental phenotypes in *N. benthamiana* and Arabidopsis leads us to propose that they may act as symptom determinants in okra; interfering with this specific effect of C4 might pave the way to engineer tolerance in this crop.

A shared property of BYVMV C4 and OELCuV C4 is their ability to suppress the spread of bleaching in SUC:SUL plants, a proxy for the intercellular spread of silencing, perhaps enabled by their targeting of BAM1 (Figure 6; Figure 7). While symptom development may predominantly affect insect vector attraction (9), impairing the advance of RNA silencing ahead of the front of infection could have a direct impact on the viral performance in the plant. It has been proposed that, to exert this effect, C4 from other geminiviruses interacts with BAM1 with particular strength at plasmodesmata, conduits for the silencing signal (17,18,21); BiFC experiments suggest that this is likely the case for BYVMV C4 and OELCuV C4 as well.

Efficient control of virus-induced diseases in crops relies on a detailed understanding of the molecular mechanisms that underlie pathogen infection and host manipulation, as such knowledge provides the foundation for the rational development of durable antiviral strategies. In this study, we identify the C4 protein encoded by the major geminiviruses threatening okra production in India, BYVMV and OELCuV, as a potentially central viral factor implicated in host-virus interactions. Our findings therefore highlight C4 as a promising candidate target for antiviral intervention. Future studies aimed at dissecting precise functions and host interactors as well as evolutionary conservation will be essential to fully evaluate the potential of C4 as target for effective and sustainable control strategies against geminivirus infections in okra.

## MATERIALS AND METHODS

### Plasmids and cloning

The BYVMV, OELCuV, and TLCYnV clones used as templates are AF241479.1, KX553923.1, and HF674921, respectively. All primers used in this study are listed in Supplementary table 1. *Escherichia coli* strain TOP10 was used for general cloning and subcloning procedures; strain DB3.1 was used to propagate Gateway-compatible empty vectors. Gateway cloning (Thermo Scientific) was followed to construct all the binary vector clones. ORFs coding for BYVMV C4, OELCuV C4, TLCYnV C4, AtRLD3, and NbSKη were amplified using the corresponding primers (Supplementary table 1) and initially cloned into pDONR/Zeo entry vector through BP reactions. LR reactions were performed to recombine them into pGWB502, pGWB505, pGWB506, pGWB554 (22), pGTQL1211YC, and pGTQL1211YN (23) to generate pGWB502-/pGWB505-/pGWB506-/pGTQL1211YC-/pGTQL1211YN-BYVMV C4, pGWB502-/pGWB505-/pGWB506-/pGTQL1211YC-/pGTQL1211YN-OELCuV C4, pGWB505-/pGTQL1211YC-/pGTQL1211YN-TLCYnV C4, pGWB554-/pGTQL1211YC-/pGTQL1211YN-AtRLD3 and pGWB554-/pGTQL1211YC-/pGTQL1211YN-NbSkη.

Y2H constructs of BYVMV C4, OELCuV C4, and TLCYnV C4 were generated by cloning in pGADT7/pGBKT7 vectors through digestion followed by ligation or by In-fusion cloning. The details of restriction enzymes used are mentioned in Supplementary table 1. PVX-BYVMV C4 and PVX-OELCuV C4 were generated through Golden Gate Assembly by *Bsa*I cut-ligation in pICH1130-PVX (Icon Genetics GmbH).

pGADT7/pGBKT7-TYLCV C4/NbSKη/AtRLD3 are described in (9). pGADT7/pGBKT7-AtBAM1 and the binary vectors to express TYLCV C4-GFP, TYLCV C4-nYFP/cYFP, AtBAM1-RFP and AtBAM1-nYFP/cYFP are described in (18), and that to express PDLP5-mcherry is described in (24).

### Plant materials

All Arabidopsis transgenic plants used in this work are in the Columbia-0 (Col-0) ecotype. The SUC:SUL transgenic line is described in (20). Plants were grown in a controlled growth room under long-day conditions (16 h light/8 h dark), with Arabidopsis maintained at 22°C and *Nicotiana benthamiana* at 25°C.

For *in vitro* culture of Arabidopsis, seeds were thoroughly surface sterilized and plated in half-strength Murashige-Skoog (½ MS) medium containing 1% sucrose and 1% agar (pH 5.7, KOH). They were stratified for 2 days at 4°C in the dark and finally grown under long-day conditions.

### *Agrobacterium*-mediated transient expression and infection assays in *N. benthamiana*

*Agrobacterium tumefaciens* strain GV3101 harbouring the corresponding constructs was liquid-cultured in Luria-Bertani (LB) medium with appropriate antibiotics at 28°C overnight. Bacterial cultures were then centrifuged at 4000 *g* for 10 min and resuspended in infiltration buffer containing 10 mM MgCl_2_, 10 mM MES (pH 5.6), and 150 μM acetosyringone. The final OD_600_ was set to 0.2-0.5. The *A. tumefaciens* suspensions were maintained in the dark for a minimum of 2 h before infiltration. The abaxial side of fully expanded young leaves of 3-4-week-old *N. benthamiana* plants were infiltrated using a 1 mL needleless syringe. For co-infiltrations, equal volumes of each culture were mixed.

PVX infections were carried out by agroinfiltration in 17-day-old *N. benthamiana* plants following the above-described protocol, with a final OD_600_ = 0.05.

### Confocal imaging

*N. benthamiana* plants were agroinfiltrated with clones to express free GFP or GFP-fused C4 proteins individually or in combination with LTI6B-RFP or PDLP5-mCherry. Samples were imaged at 2 days post-inoculation (dpi) on a Zeiss LSM880 Upright confocal laser scanning microscope using the preset settings (GFP-Ex: 488 nm, Em: 500–550 nm; RFP and mCherry-Ex: 561 nm, Em: 580–630 nm). Z-stack maximum projections were generated with ImageJ.

### Generation of transgenic plants

Wild-type and SUC:*SUL* Arabidopsis plants were transformed with pGWB502-BYVMV C4 and pGWB502-OELCuV C4 to generate 35S:BYVMV C4 and 35S:OELCuV C4, respectively, through the floral dip method (25). Empty vector (EV)-transformed Arabidopsis were also generated in parallel as control. *A. tumefaciens* strain GV3101 carrying the corresponding plasmids were grown in LB liquid medium with appropriate antibiotics at 28°C overnight. The bacterial cultures were pelleted by centrifugation at room temperature at 4000 *g* for 10 min and then resuspended in transformation solution containing 5% sucrose and 0.02% Silwet L-77. 3-4-week-old flowering plants were selected for dipping. Inflorescences were immersed in the bacterial suspension for 10–20 s with gentle agitation. Dipped plants were initially wrapped in plastic film and left in the dark for 18-24 h and then transferred to normal long-day conditions. 25 lines per construct were selected on hygromycin plates in the T1 generation; lines showing developmental phenotypes were taken to the T2 generation. 10 lines from the T2 generation were selected for further detailed characterization after confirming 3:1 segregation and assessing the transgene expression levels by qRT-PCR.

### RNA extraction and quantitative RT-PCR

For total RNA extraction, the citrate-citric acid method (26) was used. 500ng of DNase I-treated (Thermo Scientific) RNA was taken for cDNA synthesis with the PrimeScript RT MasterMix (Takara), following the manufacturer’s instructions. The subsequent qPCR reaction was performed with PowerTrack SYBR Green Mastermix, 1:100 diluted cDNA template and corresponding primers (Supplementary table 1), in a C1000 Touch Thermal Cycler (Bio-Rad) with the following program: 2 min at 95°C, and 40 cycles consisting of 15 s at 95°C, 1 min at 60°C. *AtActin* and *NbActin* were used as normalizers for qRT-PCR in Arabidopsis and *N. benthamiana*, respectively. Comparative analyses of transcripts were performed by applying the 2^−ΔCt^ method.

### Co-immunoprecipitation and protein analysis

0.8-1 g of infiltrated *N. benthamina* leaves were harvested at 2 dpi. Protein extraction, co-immunoprecipitation, and immunoblot were performed as described in (27). The primary and secondary antibodies used are the following: mouse anti-RFP (ChromoTek, 6G6, 1:5000 dilution); goat anti-GFP (SICGEN, AB0020-500, 1:5000 dilution); anti-mouse peroxidase (Sigma-Aldrich, A2554; 1:15000 dilution); anti-goat peroxidase (Sigma-Aldrich, A8919, 1:20000).

### Bimolecular fluorescence complementation

Leaf discs from *N. benthamiana* plants agroinfiltrated with clones to express the corresponding proteins were imaged at 2 dpi on a Zeiss LSM880 Upright laser scanning confocal microscope using the preset settings for YFP (Ex: 514 nm, Em: 525–575 nm).

### Yeast two-hybrid assay

The Frozen-EZ Yeast Transformation II Kit (Zymo) was used for transforming all constructs in *Saccharomyces cerevisiae* Y2H Gold strain (Clontech) following the manufacturer’s instructions. Selection of transformants and further plating were followed as described in (9).

### Quantification of *SUL* silencing

The quantification method has been previously described in (9,18). In brief, pictures of the leaves of the respective transgenic plants were converted into black and white images (8 bits). *SUL* silencing spread was quantified by calculating the total area and the black/white pixel ratio using ImageJ.

## Statistical analysis

Statistical multiple comparisons of means were performed by applying one-way ANOVA followed by Dunnett’s test (comparisons of multiple groups to one reference group). Significance level was kept at 0.05. Statistical analyses were performed in GraphPad Prism v.10 software.

## ACKNOWLEDGEMENTS

The authors thank Shaojun Pan for critical reading of the manuscript, and Bettina Stadelhofer and the central facilities at the ZMBP, especially the plant cultivation and the microscopy facilities, for excellent technical support. The authors acknowledge the financial support from the Department of Science and Technology – Science and Engineering Research Board (DST-SERB), Government of India, under the sponsored project Ref. No. EEQ/2022/000909; this work was partially funded by the DFG (TRR 356/1 2023 – 491090170) and the European Research Council (GemOmics; 101044142). AC acknowledges support from the DAAD bi-nationally supervised doctoral programme and DST-INSPIRE Fellowship (DST, Government of India).

