## Supplementary figures for "Characterization of the C4 proteins encoded by okra-infecting geminiviruses in India"

### SUPPLEMENTARY TABLES

**Supplementary table 1.** Primers used in this work.

### SUPPLEMENTARY FIGURES

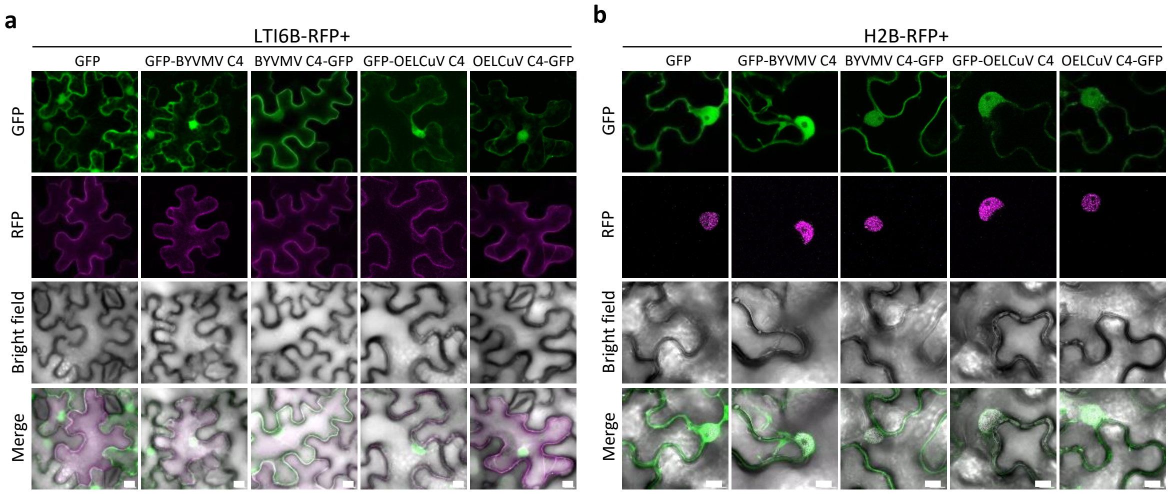


**Supplementary figure 1. Colocalization analysis of GFP-fused C4 proteins in *N. benthamiana* epidermal cells.** (a) Maximum projection of z-stack images of GFP-fused C4 proteins and the plasma membrane marker LTI6B-RFP in *N. benthamiana* leaves co-infiltrated with *A. tumefaciens* carrying constructs expressing LTI6B-RFP and GFP fused proteins, or free GFP as a negative control. Images correspond to the single-plane images in Figure 3A. (b) Colocalization analysis of GFP-fused C4 proteins and the nuclear marker H2B-RFP. *A. tumefaciens* carrying constructs to express free GFP, or GFP fused C4 proteins were infiltrated in transgenic reporter line of *N. benthamiana* stably expressing H2B-RFP. Images were taken at 2 days post-inoculation (dpi). Scale bar: 10 µm.

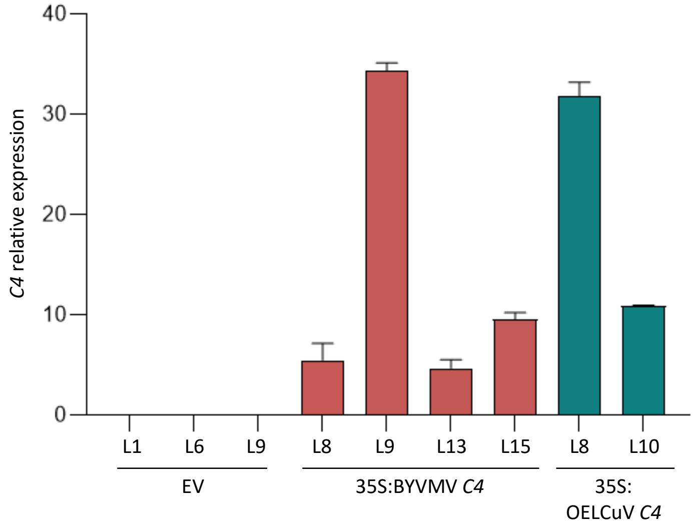


**Supplementary figure 2. C4 expression in transgenic plants.** Accumulation of BYVMV or OELCuV C4 RNA in transgenic plants relative to actin mRNA, as measured by qRT-PCR. Each bar represents the mean of five 2-week-old seedlings from each independent line. Error bars represent SEM.

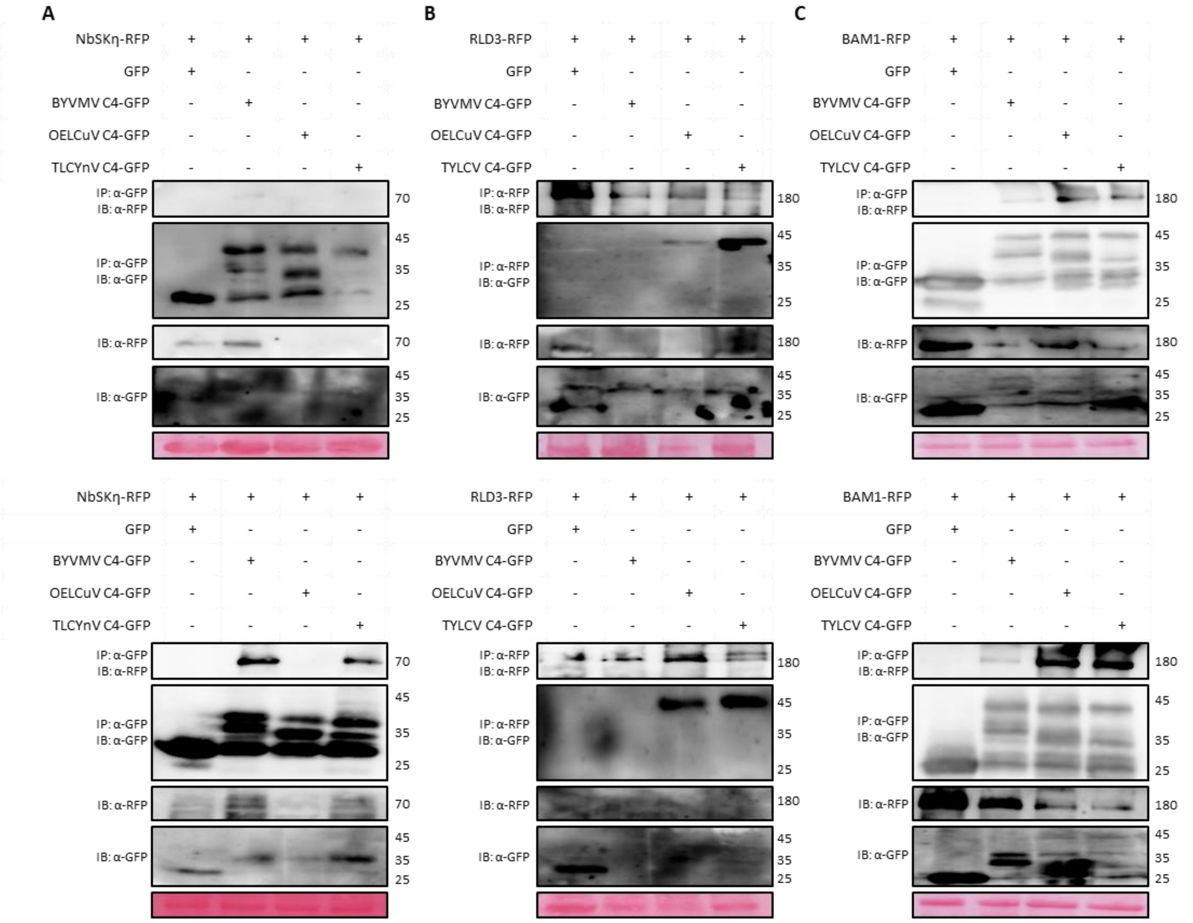


**Supplementary figure 3. Additional replicates of co-immunoprecipitation (co-IP) assays from Figures 5 and 7.** (a-c) Co-immunoprecipitation assays of GFP-fused BYVMV or OELCuV C4 with NbSKη- (a), RLD3- (b) and BAM1- RFP (c).
